# Efficient 3D Cell Tracking by Integer Programming Guided by Self-Supervised Division Scores

**DOI:** 10.64898/2026.11.17.767274

**Authors:** Tong Hu, Hironori Shigeta, Fuyu Zheng, Yuki Matsumoto, Shigeto Seno

## Abstract

Recognizing divisions in cell tracking is difficult when cells are densely packed and divide frequently. A daughter can lie closer to a neighbor than to its mother, misleading distance-based linking. Visual cues can reduce such errors, but learning them often requires manual lineage annotations, which are particularly costly for 3D data. We propose an efficient self-supervised tracker trained without manual lineage annotations, using biological priors to extract reliable divisions from its distance-based tracks as labels. Networks trained on these labels predict division scores from mother–daughter images and the mother’s temporal context to guide an integer linear program in resolving competing daughter assignments. Among the evaluated trackers, ours ranks first on one of three *C. elegans* datasets and second to a supervised tracker on the other two. It also needs less than half the training or tuning time of any compared method. Code is available at https://github.com/AnAncientMoon/Divtrack.

## 1. INTRODUCTION

Cell tracking in 3D time-lapse fluorescence images follows cells over time and identifies divisions. Many high-performing methods use tracking-by-detection, first detecting cells in each frame and then linking them across time [1]. Detection has become relatively well established, but reconstructing complete tracks and resolving divisions remain challenging [2]. An incorrect mother–daughter assignment can corrupt the inferred ancestry of all descendants. Resolving divisions correctly is therefore essential for reliable cell tracking.

Identifying cell divisions is particularly challenging when cells divide frequently and many division events occur simulta-neously or within a short time window. The *Caenorhabditis elegans* (*C. elegans*) embryo is a representative example [3]. When a cell divides, its two daughters move apart, and in a minority of divisions one of them ends up closer to a neighboring cell than to its own mother. The mother and this neighbor then compete for the same daughter, and distance-based linking favors the neighbor (Fig. 1). Dividing cells have a distinct morphology [4], so images can help identify the dividing mother cell and its daughters and distinguish them from nearby cells. Learning-based methods have achieved high accuracy in identifying divisions by learning such visual patterns from many examples, but their training often requires substantial manual annotation. Manual annotation of 3D data is time-consuming, and for large datasets it is often beyond human capacity [5]. These methods can also be computationally expensive to train on 3D image sequences.

**Fig. 1.**
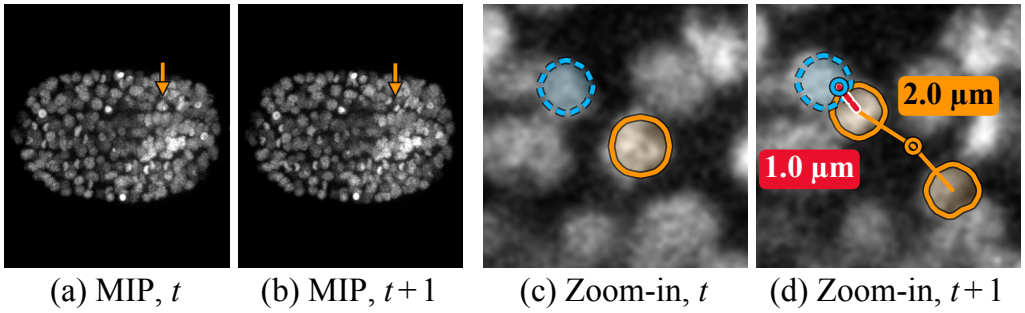
A division in a *C. elegans* embryo, shown in two consecutive frames *t* and *t* + 1. (a,b) Maximum intensity projections (MIPs) of the raw images, with arrows at the division. (c,d) Zoom-ins (maximum of two *z*-slices) with the detected mother and daughters (orange) and neighbor (dashed blue). Circles in (d) mark the mother and neighbor at *t*. Distance-based linking assigns one daughter to the neighbor (red dashed) instead of its mother (orange).

Existing methods either require manual annotations or are less reliable without them. Hirsch et al. and OrganoidTracker 2.0 (OT 2.0) use predicted cell states or division probabilities in graph optimization [4, 6], and Trackastra learns pairwise associations [1]. Unlike these three, which require manual lineage annotations, Attrackt learns associations without them by predicting each cell’s future appearance, but reports more missed and spurious divisions than with supervised training on both evaluated datasets [7]. Ultrack and PAST-FR use overlap or motion cues for linking [5, 8], which can favor the neighbor. Gilad et al. score each mother and two candidate daughters with symmetry cues to detect divisions, but can mistake similar neighboring non-dividing cells for divisions [9]. Reliable division recognition without manual annotations thus remains difficult.

To address these limitations, we propose an efficient 3D cell tracking framework that learns to recognize divisions from images without manual lineage annotations. Since distance-based linking is reliable and efficient in most cases, we retain it and introduce visual division cues to correct mother–daughter assignment errors. Moreover, cell division follows biological regularities shaped by cell-cycle regulation, so we use this prior knowledge to extract reliable training examples, which makes the learning self-supervised. On three public *C. elegans* embryo datasets, our method outper-forms the evaluated non-learning and unsupervised methods and is competitive with supervised trackers. In addition, unlike computationally intensive 3D tracking methods, our method requires only a fraction of their training or tuning time. Our contributions can be summarized as follows:

- We extract training labels using biological prior knowledge, enabling visual division cues to be learned without manual lineage annotations.
- We train networks to score candidate divisions and use them to guide mother–daughter assignment, correcting distance-based linking errors while keeping the framework lightweight.

## 2. METHOD

Our method builds initial tracks from given cell detections, extracts reliable divisions to train networks that score candidate divisions from images, and uses these scores to obtain final tracks. To build tracks across frames, we use an integer linear program (ILP), which is widely used [5, 6] in cell tracking. The ILP selects a consistent set of tracks from multiple candidate hypotheses, such as links and divisions. For each consecutive frame pair, the ILP assigns every cell in frame *t* + 1 to a link, division, or new track. We incorporate learned visual division scores into the ILP costs to guide cell assignment when distance favors a neighbor. Training requires no manual lineage annotations. Fig. 2 outlines the pipeline. We first solve the ILP with distance scores and extract reliable divisions from the resulting tracks as training labels (Secs. 2.1 and 2.2). Trained on these labels, a triplet network and a state head predict division scores. Finally, we rerun the ILP with these scores to refine cell assignments and improve tracking accuracy (Sec. 2.3).

**Fig. 2.**
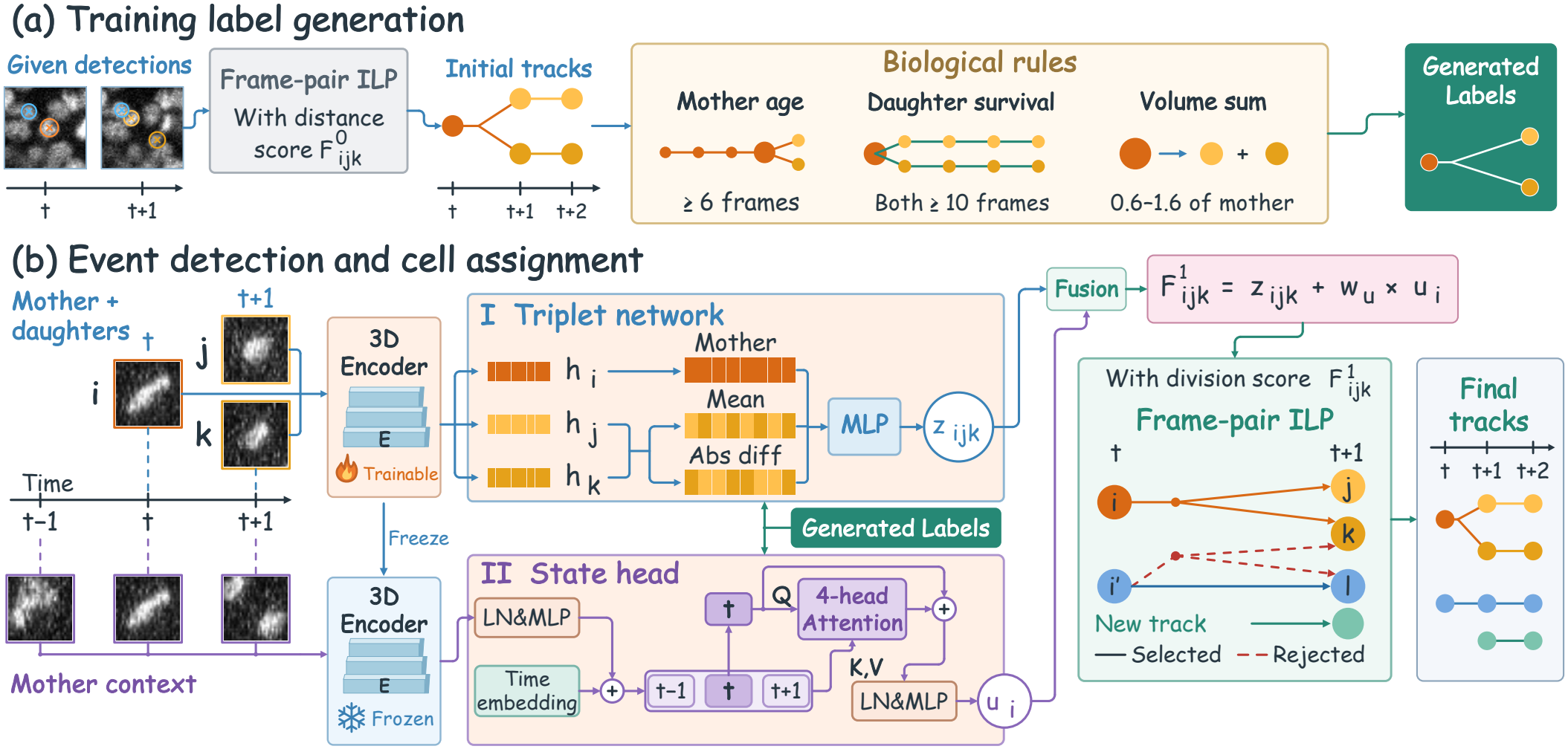
Overview. (a) The frame-pair ILP with the distance score 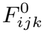 links given detections into initial tracks, from which biological rules extract divisions as generated labels. (b) The triplet network (I) scores a mother *i* with two candidate daughters *j* and *k*, and the state head (II) attends over patches at the mother’s position in frames *t −* 1, *t* and *t* + 1. Both use features from a 3D encoder *E*, trained with I and then frozen, and learn from the generated labels. Their division score 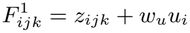 replaces 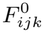 in the ILP to give the final tracks.

### 2.1. Frame-pair ILP

In cell tracking, if a division is represented by two independent pairwise links, the cost function cannot evaluate the mother and both candidate daughters together. As in the frame-pair ILP of Lou et al. [10], we instead treat a mother and two candidate daughters as one candidate division. A single variable then assigns both daughters, and a score of the whole division can enter its cost.

Let *V*_*t*_ be the set of cells in frame *t*, with *i ∈ V*_*t*_ and *j, k ∈ V*_*t*+1_ throughout. Because magnification and cell size differ between datasets, we express distance gates and sizes as multiples of the cell spacing *s*. Here, *s* is the median within-frame nearest-neighbor distance across the sequence. We only consider links and divisions between nearby cells, which rules out implausible events and keeps the ILP small. A candidate link (*i, j*) requires the distance *d*_*ij*_ between *i* and *j* to be at most 3*s*. For each cell *i* in frame *t*, we form candidate divisions (*i, j, k*) from pairs of its 4 nearest cells in frame *t* + 1, and require the two daughters to be at most 3*s* apart and each at most 2*s* from the mother. In the training sequences, these candidate gates keep 99.8% of the true divisions where all three cells are detected. Binary variables *x*_*ij*_, *x*_*ijk*_ = *x*_*ikj*_ and *n*_*j*_ select links, divisions and new tracks, at costs *c*_*ij*_, *c*_*ijk*_ and *c*_new_. We solve the following ILP:

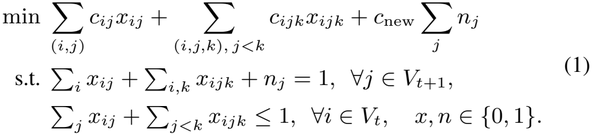

The first constraint gives each cell of frame *t* + 1 exactly one source, so at most one of two candidate divisions with a common daughter can be selected. The second lets each cell of frame *t* enter at most one link or division. The costs are

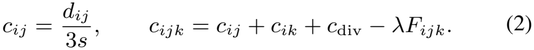

Because a division explains both daughters, its cost contains the link costs from the mother to both of them and can be compared directly with other explanations of the same daughters. In the division cost, *c*_div_ is a fixed cost paid by every division, set to 0.5 to discourage divisions without enough support. *F*_*ijk*_ scores how plausible the candidate division is, and *λ*, set to 0.03, weights it against distance, so a higher score makes the division cheaper. We set *F*_*ijk*_ to a distance score 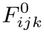 to generate training labels (Sec. 2.2) and to a division score 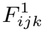 to obtain the final tracks (Sec. 2.3).

### 2.2. Generated labels

To learn the division score without manual lineage annotations, we first obtain initial tracks of the training sequences by solving the ILP with *F*_*ijk*_ set to a distance score 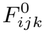. In most divisions, the two daughters lie at similar distances from the mother, and the mother is the nearest candidate mother of each daughter. We define 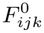 to reflect both properties,

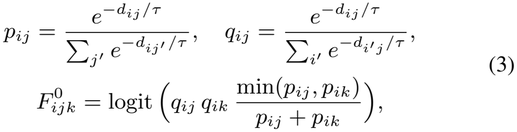

where *p*_*ij*_ is normalized over the candidate daughters *j*^*′*^ of mother *i* and *q*_*ij*_ over the candidate mothers *i*^*′*^ of daughter *j*. The temperature *τ*, set to 0.1*s*, controls how strongly *p* and *q* concentrate on the nearest candidates. The initial tracking uses *c*_new_ = 2 to favor links and divisions over new tracks and reduce missed divisions. Many initial divisions are false. Successive divisions are separated by progression through the cell cycle, and the resulting daughters typically persist as distinct cells after division. These biological expectations motivate checking the mother’s pre-division track history and the persistence of both daughter tracks. We also use mother–daughter volume consistency as a plausibility check. A division in the initial tracks is extracted as a generated label only if the mother track is at least 6 frames old at frame *t*, both daughter tracks last at least 10 frames, and their summed volume is 0.6 to 1.6 times the mother’s. Since the initial tracks rely on distance, few divisions like those shown in Fig. 1 are extracted as generated labels. Nevertheless, the division score learned from these labels can still recognize such divisions, because the appearance of dividing cells is largely independent of the distance to neighboring cells.

### 2.3. Division score

The division score combines a triplet score *z*_*ijk*_ for each candidate division with a state score *u*_*i*_ for its mother. Because a mother about to divide and its two newborn daughters look different from a non-dividing neighbor, we score the mother *i* and the two candidate daughters *j* and *k* together with a triplet network, one score for each candidate division. Its role is to identify the true division among candidate divisions that share a daughter, with the correct mother–daughter pairing receiving a higher score. The same 3D convolutional encoder *E* maps cubic patches of side 1.5*s* centered at the three cells to features *h*_*i*_, *h*_*j*_ and *h*_*k*_. Since the daughters are un-ordered, we combine their features by the mean (*h*_*j*_ + *h*_*k*_)*/*2, which summarizes their appearance, and the absolute difference |*h*_*j*_ *− h*_*k*_|, which lets the network compare the two daughters. A multilayer perceptron (MLP) with two layers takes *h*_*i*_ concatenated with these two vectors and outputs the triplet score *z*_*ijk*_ as a logit.

The triplet network is trained on the generated labels, with their divisions as positives and all other candidate divisions as negatives. As positives make up less than one in a thousand candidates, we use a binary cross-entropy loss with weighted positives. The network receives image patches but no coordinates or distances, so that it cannot directly copy the geometric decisions behind the generated labels.

The triplet network treats all candidates outside the generated labels as negatives, including missed true divisions such as those in which a mother and a neighbor compete for a daughter. We therefore add a state head that only judges whether cell *i* itself is dividing at frame *t*. In such a conflict, *u* helps tell which of the two is dividing. In frames *t −* 1, *t* and *t*+1, we crop patches as for the triplet network, all centered at the position of cell *i* in frame *t*, and extract features with the encoder *E*, which stays frozen so that this step does not change the triplet score. Each feature is projected by a shared MLP and added to a learnable time embedding for its time step. Multi-head attention uses frame *t* as the query and the three frames as keys and values, and an MLP outputs the state score *u*_*i*_ as a logit.

The state head is trained with a class-balanced binary cross-entropy loss. Its positives are the mothers in the generated labels, and its negatives are conservatively selected cells, linked reliably at every step from frame *t −* 1 to *t* + 2: the cell and its match are mutual nearest neighbors, the displacement is at most 0.5*s*, the second-nearest candidate is at least twice as far, and the volume ratio lies between 0.8 and 1.25.

The division score is a weighted sum of the two scores,

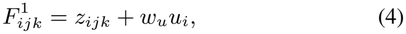

where *w*_*u*_ controls the contribution of *u* relative to *z* and is fitted by class-balanced logistic regression. Divisions in the generated labels are used as positives, while candidate divisions whose mothers are reliably linked are used as negatives. For final tracking, we set 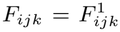 and *c*_new_ = 1, where 1 is the largest link cost within the gate. This ensures that starting a new track is never cheaper than forming a link.

## 3. EXPERIMENTS

### 3.1. Setup

We use three public datasets of *C. elegans* embryos. Of the two sequences of the confocal dataset Fluo-N3DH-CE from the Cell Tracking Challenge (CTC) [2], denoted CE below, we train on one and test on the other. Among three embryos from mskcc-confocal [11], another confocal dataset denoted mskcc, we train on two and test on the third. To test transfer to light-sheet imaging, all methods apply their trained models and the parameters selected on mskcc to three nih-ls embryos [12, 13] without further training or tuning, using their first 270 frames before twitching begins. As in the Cell Linking Bench-mark (CLB) of the CTC, all methods work on the same given detections: the CLB masks (ERR SEG) for CE, and for the other two datasets, spheres at the annotated cell positions with 0.8 times the annotated radius to reduce overlaps.

The compared methods are Ultrack [5], PAST-FR [8], Trackastra [1], OT 2.0 [4], Hirsch et al. [6] and Attrackt [7], grouped in Table 1 by the supervision they require. All methods use the same training and test splits. Methods that train networks, ours included, train one model on the three pooled training sequences. Our results are averaged over five random seeds, whereas each compared method is run once. For Hirsch et al., we use their cell-state classifier and ILP, tuning the ILP weights on the ground truth, and omit their 4D U-Net detector since detections are given.

**Table 1.** Results on three *C. elegans* embryo datasets with given detections. Scores on nih-ls are averaged over its three embryos.

|  | CE |  | mskcc |  | nih-ls |  | Time |
|---|---|---|---|---|---|---|---|
| Method | OP<sub>CLB</sub> | BC(1) | OP<sub>CLB</sub> | BC(1) | OP<sub>CLB</sub> | BC(1) | (h) |
| *No network training, parameters tuned on training ground truth* |  |  |  |  |  |  |  |
| Ultrack | 0.813 | 0.569 | 0.895 | 0.768 | 0.892 | 0.733 | 8.0 |
| PAST-FR | 0.931 | 0.865 | 0.955 | 0.900 | 0.935 | 0.844 | 5.6 |
| *Trained on manual lineage annotations* |  |  |  |  |  |  |  |
| Trackastra | 0.917 | 0.883 | 0.955 | 0.926 | 0.967 | 0.921 | 8.2 |
| OT 2.0 | 0.934 | 0.919 | 0.976 | 0.966 | 0.969 | 0.935 | 9.9 |
| Hirsch et al. | 0.935 | 0.909 | **0.992** | **0.992** | **0.984** | **0.968** | 10.1 |
| *No manual lineage annotations* |  |  |  |  |  |  |  |
| Attrackt | 0.905 | 0.849 | 0.980 | 0.967 | 0.968 | 0.922 | 23.6 |
| Ours | **0.952** | **0.926** | 0.983 | 0.970 | 0.977 | 0.944 | **2.3** |

To evaluate overall tracking performance, we report OP_CLB_ [14], the mean of two CTC measures: LNK, which scores how well cells are followed across frames, and BIO, which averages biological measures of complete tracks, track fractions, cell-cycle lengths and divisions. We compute BIO with BC(1), the division F1 score requiring matched mother–daughter branches with a one-frame timing tolerance, and also report BC(1) separately. All times in Table 1 are hours of training or tuning for one seed, measured on a machine with one RTX PRO 6000 GPU and 8 CPU cores.

### 3.2. Results

Without manual lineage annotations, our method achieves the highest OP_CLB_ and BC(1) on CE. On mskcc and, without retraining, on light-sheet nih-ls, it is second only to Hirsch et al., who use ground truth for both training and parameter selection. Its training, 2.3 hours per seed, takes less than half the training or tuning time of any compared method. We also submitted our method to the CTC, whose organizers evaluated it on the hidden test sequences of the CLB, where it achieved state-of-the-art performance on Fluo-N3DH-CE with an OP_CLB_ of 0.950.

### 3.3. Analysis

On CE and mskcc, the distance baseline (*F*_*ijk*_ = 0) misses 99% and 96% of divisions where a daughter is closer to a neighbor than to its mother (Fig. 1), usually predicting a false division of the neighbor, whereas our method misses only 24% and 14%.

To understand the triplet network’s decision, we compare the true division (*i, j, k*) with the false division (*i*^*′*^, *k, l*) favored by distance (Fig. 3a). Both contain daughter *k*, which the false division assigns to neighbor *i*^*′*^ alongside its continuation *l*. Fig. 3b shows two cosine similarities computed from the triplet network’s encoder features, one between the two daughters and one between each mother and the mean of other true mothers. Both are generally higher for true divisions, so the features tell them apart where distance fails. To compare mother and daughter contributions to *z*, we replace the false division’s mother and non-shared daughter with those of the true division, in either order. A two-factor Shapley decomposition [15] averages each replacement’s effect over the two orders, separating the score difference into mother and daughter contributions. The daughter contribution accounts for 87% on CE and 62% on mskcc. Daughter features thus contribute most to the score difference. Un-like Gilad et al.’s hand-crafted symmetry cue [9], these features are learned from generated labels.

**Fig. 3.**
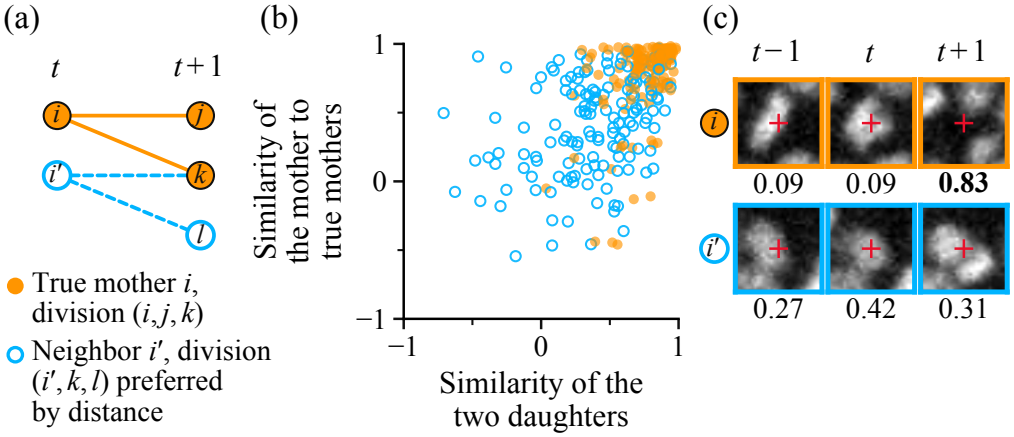
Analysis of divisions of the kind shown in Fig. 1. (a) Compared cells. (b) Cosine similarities of encoder features. (c) Middle slices of the state head’s patches at *i* and *i*^*′*^ of Fig. 1 (cross), with each frame’s attention. Values are means of five seeds.

The state head separately judges whether the mother itself is dividing. Replacing its *t* + 1 features with those of another cell lowers *u* by 3.7 on CE and 5.3 on mskcc on average, whereas replacing *t −* 1 features has almost no effect. Attention also favors *t* + 1 far more for true mothers than for neighbors (Fig. 3c). At *t* + 1, daughters have moved apart, often leaving the mother’s former position empty, whereas the neighbor remains. Across all five seeds, daughter features contribute most to the triplet score difference between true and false divisions, while the state head assesses temporal changes at the mother’s position. These complementary visual cues help correct distance-based assignment errors.

### 3.4. Ablation

To measure our gain over the distance baseline in Table 2, we remove both scores, lowering OP_CLB_ by 0.072 on CE and 0.039 on mskcc. Removing one score at a time tests whether both are needed: either score alone recovers most of this loss, but removing either lowers OP_CLB_ in every seed, except removing *z* on mskcc. There, removing *z* sends more daughters to neighbors but misses fewer divisions in crowded frames, probably because the generated labels miss many divisions there, which the triplet network learns to reject. To test whether the triplet score’s gain comes from each candidate’s own image, we shuffle patches among each frame’s candidates during training, leaving labels in place. This keeps each frame’s images and label proportions, yet OP_CLB_ falls to or below the distance base-line, so the triplet network relies on each candidate’s own image.

**Table 2.**
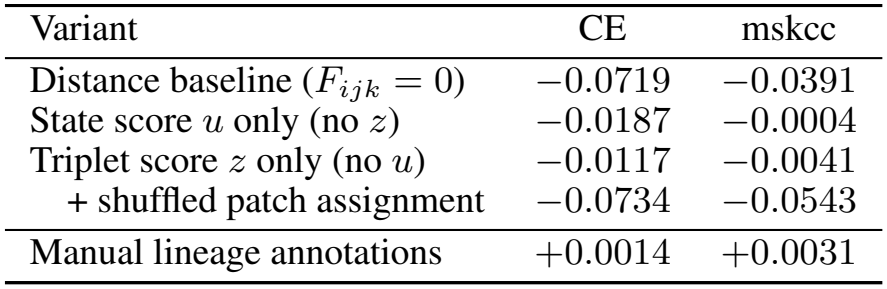
Ablation. Difference in OP_CLB_ from the full model with the same seed, averaged over five seeds (three for patch shuffling). The indented row changes only the training of the triplet-only model.

To measure the cost of using generated labels, we train both networks on labels from the manual lineage annotations, which are more complete. This raises OP_CLB_ by only 0.001 on CE and 0.003 on mskcc, so generated labels cost little in tracking performance.

## 4. CONCLUSION

We presented a 3D cell tracker in which a frame-pair ILP links cells, and self-supervised networks trained on generated labels from its first pass supply visual evidence for division costs. Without manual lineage annotations, it recovers most divisions on CE and mskcc in which distance assigns a daughter to a neighbor, and it is competitive with supervised methods on three *C. elegans* datasets with less than half their training time. Our evaluation covers only *C. elegans* embryos with fluorescently labeled nuclei and given detections, and future work will extend it to other species and to automatic detection.

## 5. COMPLIANCE WITH ETHICAL STANDARDS

This research study was conducted retrospectively using images of *C. elegans* embryos made available in open access. Fluo-N3DH-CE is used with the permission of the Cell Tracking Challenge organizers, and mskcc-confocal and nih-ls are released under the CC BY 4.0 license. No experiments on human subjects or animals were performed for this study, and ethical approval was not required.

## 6. ACKNOWLEDGMENTS

This work was supported by JSPS KAKENHI Grant Number JP26H02546. The authors have no relevant financial or non-financial interests to disclose. Claude was used to assist with manuscript preparation and language polishing in each section, and with code generation for the experiments. The authors reviewed and verified the final content and take full responsibility for the work.

